# 3-D navigation – A core driver of brain plasticity in video game interventions

**DOI:** 10.1101/453613

**Authors:** Simone Kühn, Maxi Becker, Jürgen Gallinat

**Author notes:** Corresponding author: Simone Kühn.

## Abstract

Recent evidence has repeatedly shown that 3D platform video game training leads to substantial brain structural plasticity in hippocampus, prefrontal cortex and cerebellum. However, a great disadvantage of using complex video game interventions is the difficulty to attribute the observed effects to specific game mechanics.

In order to address this caveat, we conducted a longitudinal training study in which 40 participants were randomly assigned to train with a 3D platformer game or a 2D platformer game. The main difference between the two games lies within their affordance for spatial exploration. After a training phase of two months, we observed extended brain structural increases in the 3D in comparison to the 2D condition in bilateral prefrontal areas, hippocampus/ entorhinal cortex as well as precuneus and the temporal lobe. In the reverse contrast an increase in bilateral caudate nucleus was observed.

The results demonstrate a crucial role of 3D spatial navigation for widespread brain plasticity effects within a two-months training setup. Given the vast complexity of video games, spatial navigation seems to play an outstanding role in structural plasticity. Since prefrontal, temporal and hippocampal volume deficits are prominent risk factors for several psychiatric disorders, daily navigation habits (outdoor movement, using GPS devices etc.) have to be considered in future mental disorders prevention research.

---

In the past years scientific evidence has accumulated indicating that video game playing may have a positive impact on brain structure and function. This evidence is of great societal relevance since video gaming is becoming more and more pervasive across all age groups. It has been estimated that people spend a collective three billion hours per week playing video gamesworld-wide^1^, illustrating the importance of research investigating the relationship between video game play and brain plasticity. First, cross-sectional evidence has hinted at differences between gaming genres in their effects on brain structure; with Logic and Puzzle game play (e.g. *Tetris*) and platform game play (e.g. *Super Mario64*) being positively associated with grey matter volume in the entorhinal cortex, a region strongly linked to the hippocampus (HC),while action role-playing game play (e.g. *Borderlands 2*) being negatively associated ^2^. However, cross-sectional evidence leaves it impossible to determine whether observed brain correlates of gaming are due to pre-existing differences in the brain or whether they represent the results of intense video-game training. Here, longitudinal interventional research contributes causal knowledge on the directionality of the effects. We were able to demonstrate that two months of training (30 min/day) with the 3D platform game (*Super Mario 64*) elicited extensive brain structural plasticity effects (gray matter volume increase) in right HC, right dorsolateral prefrontal cortex (DLPFC) and cerebellum in young healthy adults ^3^. The hippocampal and cerebellar plasticity effects of the same 3D platform game have been replicated in a study on older adults that have been trained for six months ^4^. And yet another recent study has shown brain plasticity effects in HC/entorhinal cortex in response to training with 3D platform games in a younger sample^5^. Interestingly, the observed hippocampal volume increases for 3D platform games contrasted with hippocampal volume decreases elicited by action game play in participants using non-spatial navigation strategies. These opposing effects elicited by the two different game genres point at the need to unravel the game mechanics of video games that elicit brain plasticity. Unfortunately, the search for these game mechanics is difficult when off-the-shelf video games are used for training, since they usually comprise a complex set of mechanics that make it difficult to attribute observed effects (e.g. brain structure and cognition) onto a single feature of the game. On the other hand the great appeal of using off-the-self video games as training intervention tools is that they are designed to capture players’ attention, maintain an enjoyable level of challenge and provide a sense of accomplishment ^6^. The importance of these attributes is reflected by empirical data showing a positive association between the self-reported fun while playing the game with brain plasticity effects elicited ^3^.

However, the present literature offers first hints at the potential game mechanics within the 3D version of Super Mario. In the first study reporting HC plasticity in response to 3D Super Mario playing, these effects were associated to changes in navigation strategy; that is participants showing stronger grey matter volume changes in HC likewise displayed a shift from an egocentric to an allocentric navigation strategy^3^. This finding is in line with previous research showing that allocentric navigation strategy relies on hippocampal activation ^7, 8^. Additionally, a behavioural study focussed on changes in cognitive performance after two weeks of training with 3D Super Mario as compared to a 2D game (*Angry Birds*), demonstrating increases in spatial performance in a virtual water maze and in episodic memory, both tasks covering domains that have been associated to hippocampus integrity ^9^. Therewith, both studies suggest spatial processing as a candidate mechanism for plasticity as well as relevant cognitive transfer effects.

Within the present study, we set out to demonstrate that the brain plasticity effects observed in HC ^3-5^, as well as in DLPFC ^3^ are elicited by the training of spatial processing within a 3D environment and are not present in a 2D game in which spatial processing is limited. To test this hypothesis, we examined brain plasticity changes after two months of daily video gaming training with the 3D platformer *Super Mario 64* and compared it to changes elicited after the same amount of game play in a 2D side-scroller version *Super Mario Bros*.

## Methods

### Participants

The local ethics committee of the Medical Association, Hamburg, Germany, approved of the study. Forty healthy participants (mean age = 27.4, SD = 6.6, 18 females) were recruited by means of newspaper and internet advertisements. After complete description of the study, the participants’ informed written consent was obtained. According to personal interviews (Mini-International Neuropsychiatric Interview) participants were free of mental disorders. In addition, exclusion criteria for all participants were abnormalities in MRI, relevant general medical disorders and neurological diseases. Participants reported little, preferably no video-game usage in the past year (none of the participants ever played neither *Super Mario 64* nor *Super Mario Bros*. series intensely in their lifetime). The participants received a financial compensation for the testing sessions, but not for the video game training itself.

### Training Procedure

The participants were randomly assigned to the 3D experimental group or the 2D active control group. The 3D group (n=20, mean age = 28.8, SD = 6.8 years, 7 females, years of education = 16.6) was instructed to play the video game *Super Mario 64* on a portable Nintendo Dual Screen (DS) XXL console for at least 30 min per day over a period of two months. The 2D active control group (n=20, mean age = 26.1 years, SD = 6.4, 11 females, years of education = 16.7) received the same instruction and console but played the 2D video game *Super Mario Bros*.

*Super Mario 64* is a 3D platformer game in which a princess has to be saved. The player needs to explore the virtual environment to collect stars by exploring the levels precisely (along the x, y and z coordinate), solving puzzle or defeating enemies to be able to proceed to higher levels. The environments are full of proximal and distal landmarks and provide multiple paths to the same location, therefore players need a rich internal representation of the environment to progress in the game. On the top half of the screen the environment is seen from a third person perspective (behind the character), on the bottom half of the screen a map is shown from a bird’s eye view, enabling orientation and in particular the localization of stars (see *Figure 1*). *Super Mario Bros* is a 2D platformer game with a similar goal and reward structure as in *Super Mario 64*. The main difference is that the character can only be moved in a 2D world (along the x and y coordinate), that is designed mostly for movements from the left to the right of the screen mostly, while the side-scrolling option is used to visualize a larger world.

**Figure 1:**
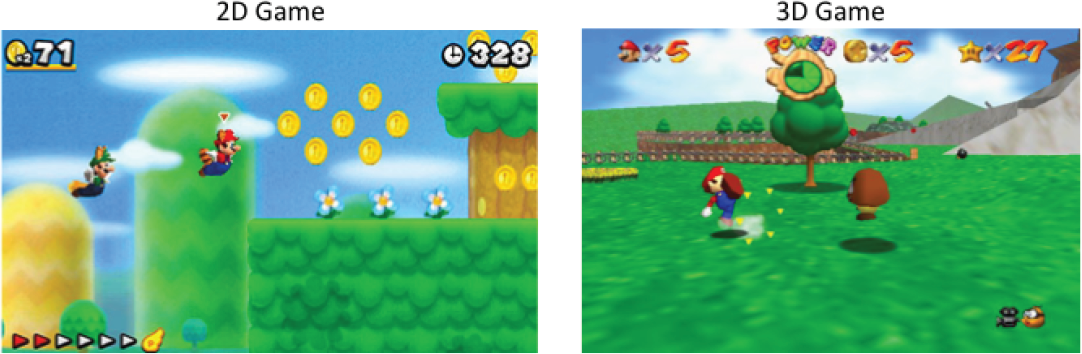
Screenshots from the two video games used in the training intervention.

Participants were instructed on how to use the keys on the gaming console and learned about the rules of Super Mario prior to the training phase. Each and every week participants filled in a questionnaire online asking about the duration of game play in the past week. Participants were instructed to play about 30 min per day over the course of two months.

### Scanning Procedure

Structural images were collected on a Siemens Skyra 3T scanner (Erlangen, Germany) and a standard 32-channel head coil was used. The structural images were obtained using a three-dimensional T1-weighted magnetization prepared gradient-echo sequence (MPRAGE) (repetition time = 2500 ms; echo time = 2.12ms; TI = 1100 ms, acquisition matrix = 240 × 241 × 194, flip angle = 9°; 0.8 x 0.8 x 0.94 mm voxel size).

### Tunnel task

Before and after the training procedure the training participants underwent the so-called tunnel task to assess navigation preferences ^3, 10^. This task was used, since previous reports on video game plasticity showed differences with navigation strategy ^5^. Participants saw a sparse visual flow, at the end of the visual flow they had to indicate the direction where the starting position was. To determine whether participants are so called “turners” (egocentric frame, participants react as if they had taken on the new orientation during turns of the path by mentally rotating their sagittal axis) or “nonturners” (allocentric frame, participants tracked the new orientation without adopting it) or rather to which degree they adopt which strategy. At the end of each path participants had to choose between two homing vectors indicating their end position relative to the origin, one being the correct answer from an egocentric, one from an allocentric perspective. To determine the frame participants use habitually, 10 trials were administered and the egocentricity ratio was computed as the fraction of egocentric choices (ego/ego+allocentric-ratio).

### Data Analysis

We obtained grey matter volume estimates using CAT12 (v1278) running on SPM12 and Matlab R2016b using default parameters. Longitudinal processing was performed using default parameters according to the standard protocol (http://www.neuro.uni-jena.de/cat12/CAT12-Manual.pdf). CAT12 automatically performs intra-subject realignment, bias correction, segmentation, and normalization (normalization is estimated for the mean image of all time points and then applied to all images). Segmentation and normalization were done with default parameters segmenting into three voxel classes (grey matter (GM), white matter (WM), and cerebrospinal fluid (CSF)) using adaptive maximum a posteriori segmentation and partial volume segmentation. The extracted GM maps were smoothed using an 8mm FWHM kernel.

Processing included several stages of quality checking: Images were visually inspected for artefacts prior to processing. Then, a statistical quality control based on inter-subject homogeneity after segmentation was conducted using the “check homogeneity” function in CAT12. The images were then visually inspected again after preprocessing.

Statistical analyses were carried out by means of a whole brain flexible factorial design with a focus on the interaction of time (pre vs. post) x group (3D vs. 2D). We tested the following contrasts: -1 -1 -1 3 and 1 1 1 -3 (2Dpre 2Dpost 3Dpre 3Dpost) since we were interested in brain regions that display a significant increase or significant decrease in the 3D compared with the 2D condition over training time. The resulting maps were thresholded using family wise error correction (FWE) *p* < 0.05 together with a non-stationary smoothness correction^11^. To prevent missing relevant information by thresholding to conservatively, we additionally employed a cluster thresholding together with non-stationary smoothness correction implemented in CAT12. To investigate the relationship between caudate and hippocampus volume changes we extracted the data from the significant clusters, subtracted pretest from posttest values and correlated the difference scores using Pearson’s correlation analysis.

To explore potential relationships between navigation strategy (egocentric vs. allocentric) and brain structural plasticity in the clusters showing a significant group x time interaction, we computed independent t-tests to compare egocentric with allocentric participants as well as correlated brain change to the egocentricity ratio (ego/ego+allocentric-ratio).

## Results

The groups did not differ in terms of age (*t*=-1.30, *p*=0.202), sex distribution (χ^2^=1.62, *p*=0.20) or education years (*t*=0.06, *p*=0.95)

On average, participants in the 3D group played 3 hours and 34 min per week (SD = 29 min) while the 2D group played 3 hours and 48 min per week (SD = 47 min) according to weekly self-reports amounting to about 30 min per day in each group. In the 3D group 13 and the 2D group 12 participants showed a preference for egocentric navigation strategy assessed via their behaviour in the tunnel task at pretest.

Since we were mostly interested in detecting regions that show different increases or decreases over time between the two training groups, we computed whole brain interactions first. We found significant increases in the 3D compared with the 2D condition over time in right DLPFC, rostral cingulate zone/ mid cingulum, left middle frontal gyrus, right precentral gyrus, left inferior frontal gyrus, right precuneus, left middle temporal gyrus/temporoparietal junction, right insula/inferior frontal gyrus (FWE *p*<0.05) and right hippocampus/ entorhinal cortex when using a more lenient threshold (*p*<0.001, cluster extent corrected) (*Table 1*, *Figure 2*). In the reverse contrast, we observed a reduction in bilateral caudate nucleus over time in the 3D vs. 2D condition when using a more lenient threshold (*p*<0.001, cluster extent corrected) (*Table 2*, *Figure 2*). Based on the previous observations that caudate nucleus and hippocampus grey matter show an inverse relationship according to the individuals’ applied navigation strategy, we tested for associations of both structures’ volume. Contrary to previous reports, we observed a positive correlation between our caudate and hippocampal cluster at both time points (Pretest: *r*(40)=0.33, *p*<0.05, Posttest: *r*(40)=0.41, *p*<0.01, similar results were obtained when controlling for intracranial volume). Additionally, we computed the correlation between grey matter volume changes over time in the two clusters that were found significant in the group x time interaction. Here, we observed a significant negative correlation between caudate nucleus and hippocampus change when considering both groups together (*r*(40)=-0.44, *p*<0.01), the 3D group only (*r*(20)=-0.46, *p*<0.05) but not the 2D group only (*r*(20)=-0.31, *p*=0.18), illustrating that higher increases in the hippocampus were accompanied by higher decreases in caudate volume. We did not find any significant differences in structural brain changes of caudate or hippocampus between participants who preferred an egocentric or an allocentric navigation strategy (both groups: *p*>0.55, 2D only: *p*>0.68, 3D only: *p*>0.48), likewise we found no significant correlations between egocentricity ratio and structural changes (both groups: *p*>0.48, 2D only: *p*>0.13 for caudate nucleus, 3D only: *p*>0.35) and no association between changes in egocentricity ratio and structural change (both groups *p*>0.59, 2D only: *p*>0.60, 3D only: *p*>0.59).

**Figure 2:**
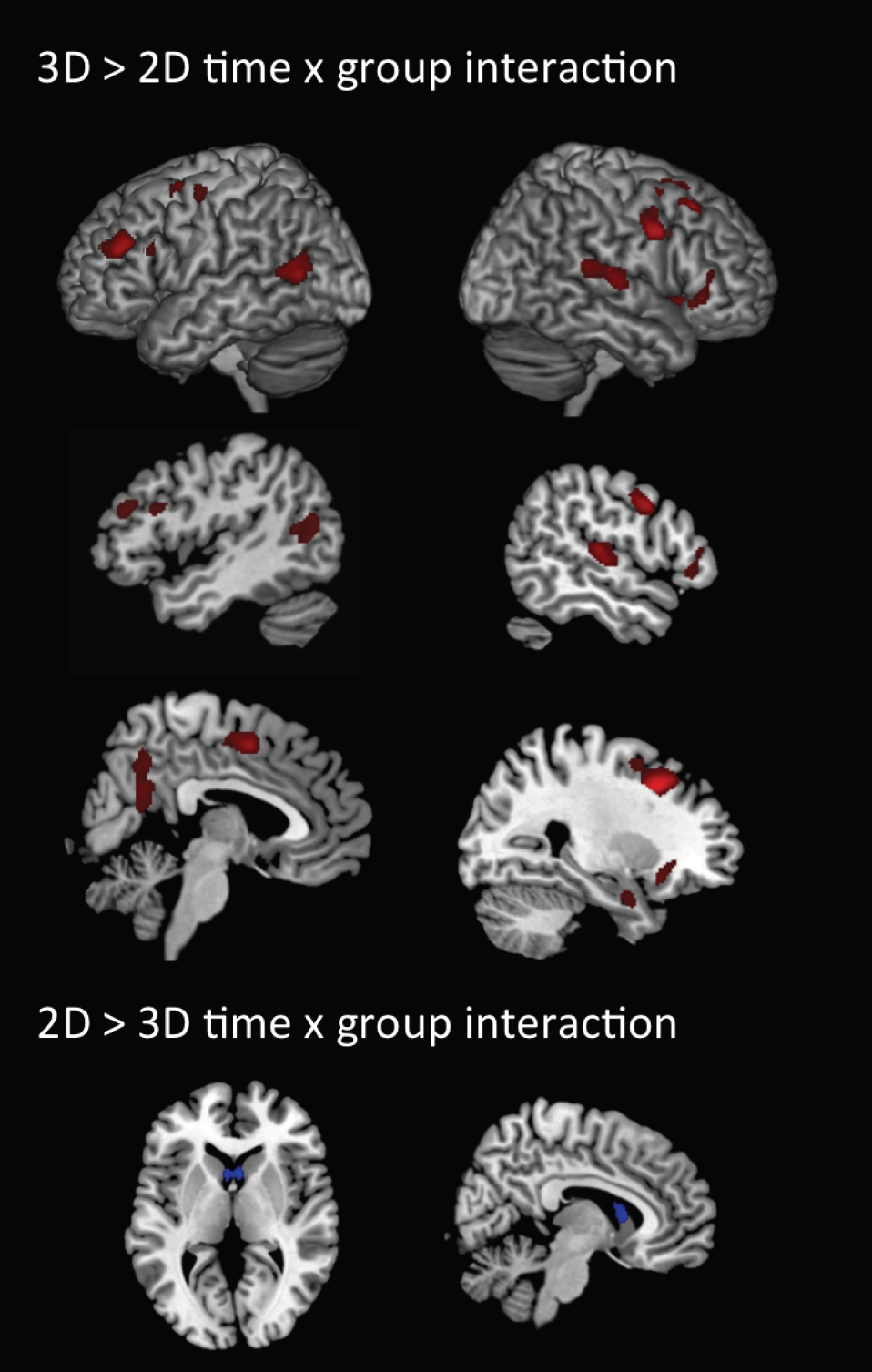
Brain regions showing a significant group (3D vs. 2D) x time (pre vs. post test) interaction in grey matter volume (all clusters are thresholded at *p* < 0.05, family wise error and non-stationary smoothness corrected, except for caudate and hippocampus which are the result of a thresholding with p<0.001, cluster extent corrected and non-stationary smoothness corrected).

**Table 1:**
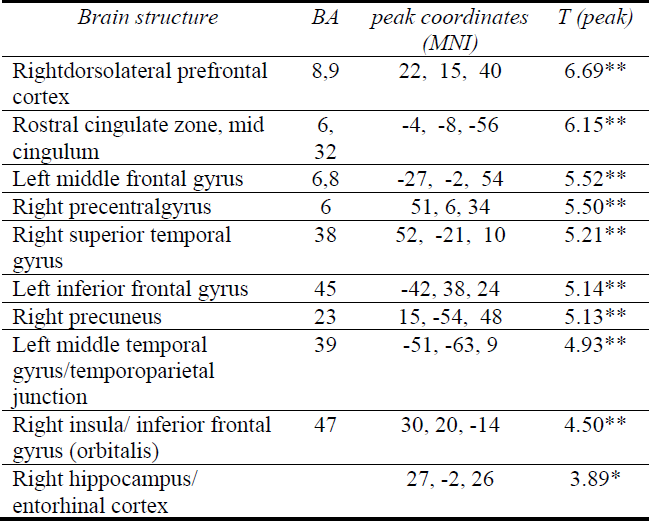
Brain regions showing a significant interaction effect of group (3D>2D) and time (pretest vs. posttest)in grey matter volume (\*\**p* < 0.05, family wise error and non-stationary smoothness corrected, * p<0.001, cluster extent corrected and non-stationary smoothness corrected).

**Table 2:**
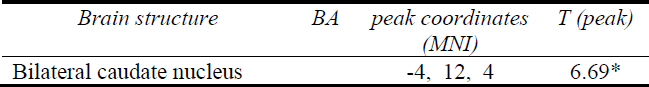
Brain regions showing a significant interaction effect of group (2D > 3D) and time (pretest vs. posttest) in grey matter volume (* p<0.001, cluster extent corrected and non-stationary smoothness corrected).

## Discussion

Within the scope of the present study we set out to unravel the crucial elements of the game that cause the structural brain plasticity observed in response to 3D video game playing. Previous studies demonstrated that playing the 3D platformer game *Super Mario 64* over a period of two to six months resulted in structural increases in the HC (in younger ^3, 5^ and older populations ^4^) as well as in right DLPFC ^3^. First evidence hints at the importance of the spatial exploration game mechanics, namely a correlation between brain plasticity in HC and changes in navigation strategy due to training ^3^ as well as behavioural effects of 3D game training on spatial navigation ability ^9^. In order to test this hypothesis experimentally, we compared the brain plasticity effects elicited by playing the 3D game *Super Mario 64* to those elicited by playing a 2D version of the game. The 3D platformer game affords that players explore a virtual environment along three dimensions (x, y and z coordinate) and are offered a map view for orientation while the 2D version of the game mostly affords a left to right movement along only two dimensions (x and y coordinate).

When comparing GM volume changes between pretest before and posttest after two months of training between the 3D and 2D training group we observed significant interaction effects with the 3D group showing increases whereas the 2D group was relatively stable in bilateral prefrontal cortex (right DLPFC, rostral cingulate zone/ mid cingulum, left middle frontal gyrus, left inferior frontal gyrus) as well as right precentral gyrus, right precuneus, left middle temporal gyrus/temporoparietal junction, right insula/inferior frontal gyrus at a very conservative threshold and additionally right hippocampus when using a more lenient threshold. Interaction effects with the opposite direction were only observed at a more lenient threshold in bilateral caudate nucleus.

### Prefrontal cortex & HC

Most interestingly, the present study revealed brain structural increases in comparable regions that we previously reported when contrasting plasticity effects elicited by 3D video game play against a passive control group ^3^, namely in right DLPFC and right hippocampus. The DLPFC cluster from the previous study shows overlap with the cluster detected in right inferior frontal gyrus/insula but the cluster in DLPFC in the present study was located slightly more ventral. The locations in the hippocampus do not overlap. The previous study comparing 3D game play against a passive control group revealed structural changes in the right body and tail of the hippocampus extending into parahippocampus, whereas the present hippocampal cluster is located in the right head of the hippocampus and the neighbouring entorhinal cortex. There is substantial evidence supporting that the posterior part of the hippocampus, namely body and tail, are involved in cognitive processing such as memory and navigation ^12^, whereas the anterior portion (namely the head) of the hippocampus modulates affective processing^13, 14^. However in rats evidence shows that the ventral hippocampus is likewise involved in the retrieval of spatial information ^15^. Moreover, the anterior hippocampal region displaying more structural plasticity in the 3D compared with the 2D group is close to the locations where West and colleagues who trained spatial learners with a 3D video game or an action video game reported structural increases as well ^5^.

### Motor-related brain regions

Within the present study we observed brain structural increases in 3D and stability in the 2D group in rostral cingulate zone and right precentral cortex. Rostral cingulate zone has been associated with the process of deciding between different response alternatives ^16, 17^, which is indeed a frequent element in video gaming. It could be that the spatial exploration in a 3D space, that is strongly linked to moving the game avatar to these locations, affords more rostral cingulate zone activation, and therewith elicits more structural plasticity than the more simple type of navigation in the 2D game version. Interestingly, the cluster in precentral cortex was located closer to the motor representation of the eye than of the hands, suggesting that the 3D version may have asked for more movement of the eye as compared to the 2D version, which may be due to the fact that the 3D version afforded eye movement between the top and bottom screen depicting the map. As expected, we did not observe any differences in cerebellar brain structural changes as in the previous *Super Mario 64* study, where we compared against a passive control group, most likely since both games afford as similar degree of fine-motor tuning of the hands while using the game controllers.

### Other brain regions

Above and beyond prefrontal, hippocampal and motor-associated brain regions, we observed significant plasticity effects in 3D vs. 2D in precuneus and left middle temporal gyrus/temporoparietal junction. Precuneus has previously been associated with training-related structural plasticity effects in younger adults in a study in which participants walked on a treadmill and navigated in a virtual zoo for four months ^18^. The cluster in middle temporal gyrus located within the so-called temporoparietal junction is not typically related to spatial processing, but due to its location at the confluence of diverse information streams it is known as a critical hub for multisensory processing ^19^, which may indeed be more relevant in a 3D compared with a 2D game context.

### Caudate nucleus and hippocampus in navigation

We observed a significant decrease in grey matter volume within bilateral caudate nucleus in the 3D compared with the 2D condition. This is remarkable since both caudate nucleus as well as the hippocampus have been shown to be differentially involved in spatial processing. Participants using spatial landmarks to navigate show increases in brain activation of the (preferentially) right hippocampus whereas participants who used a non-spatial response strategy (memorizing right and left turns from a given start position) showed increased activity in caudate nucleus during navigation ^20, 21^. This division of labour has previously been reflected in an inverse relationship of grey matter volume in hippocampus and caudate nucleus. Increased grey matter in caudate has been reported to be associated with lower grey matter volume in hippocampus in humans ^22^ as well as in animals ^23^. We were not able to replicate this finding on the clusters observed in the present study. However, we found an inverse correlation between the observed changes in brain structure over time in caudate nucleus and hippocampus. That is, the more the hippocampal grey matter volume increased, the more the caudate cluster decreased in grey matter volume. This effect was significant in the 3D group and also when taking both groups together.

Since previous research has commonly dissociated spatial strategies with hippocampal and non-spatial response strategies with caudate function and structure, we split our sample up into participants showing a preference for an egocentric or an allocentric reference frame in navigation. Although both the ego- and allocentric strategy are spatial, we had previously observed an association between hippocampal plasticity and changes towards and allocentric navigation strategy preference so that we thought there could potentially be differences between participants with different strategies, but found no evidence for this. Neither did we replicate the previous association between strategy changes and brain changes nor did we find any differences in structural changes between participants showing a habitual tendency for an egocentric or allocentric navigation strategy.

In our previous Super Mario study, we speculated that it may be the integration of the two perspectives, the third person view on the top screen and the bird’s eye view on the bottom screen that caused the observed plasticity effects. However, the studies replicating plasticity effects of Super Mario ^4, 5^ had participants play on a Wii console where the bird’s eye view is not available and still observed the hippocampus/entorhinal effects. Therefore, the hippocampal effects cannot be dependent on the integration of the two perspectives only. However, the structural increases in prefrontal cortex, that was not observed in these later studies ^4, 5^, may potentially be explained by this integration affordance rather than the exploration in the 3D environment. Therefore, further research into the game mechanics that contribute to structural brain plasticity may compare the presence of absence of the bird’s eye view on the bottom of the screen onto brain plasticity in prefrontal cortex.

The present results clearly demonstrate that the brain plasticity effects within the hippocampal formation elicited by two months of playing the 3D platformer *Super Mario 64* is based on the higher spatial exploration affordances in comparison with the 2D platformer *Super Mario Bros*. The prefrontal effects observed could potentially be caused by the above-mentioned difference in spatial exploration or the presence of a bird’s eye view that affords an integration of two different spatial perspectives. Knowing the game mechanics causing hippocampal effects is crucial in order to inform the design of targeted interventions, most importantly since hippocampal volume is an important biomarker for numerous neurological and psychiatric disorders including Alzheimer’s disease ^24^, post-traumatic stress disorder ^25^ and schizophrenia ^26^.

## Conflict of interest

The authors have no conflict of interest.

## Acknowledgement

We want to thank Wyn Weinrich, Jucia Yogendran, Ali Parrand and Andreas Hensel for acquiring the behavioral data.

The study has been funded by a grant of the German Science Foundation (SFB 936/C7).

SK has been funded by two grants from the German Science Foundation (DFG KU 3322/1-1), the European Union (ERC-2016-StG-Self-Control-677804) and a Fellowship from the Jacobs Foundation (JRF 2016-2018).

